# Identification of combinatorial and singular genomic signatures of host adaptation in influenza A H1N1 and H3N2 subtypes

**DOI:** 10.1101/044909

**Authors:** Zeeshan Khaliq, Mikael Leijon, Sándor Belák, Jan Komorowski

**Author notes:** Corresponding Author, Email Addresses: ZK ML SB JK.

## Abstract

**Background:** The underlying strategies used by influenza A viruses (IAVs) to adapt to new hosts while crossing the species barrier are complex and yet to be understood completely. Several studies have been published identifying singular genomic signatures that indicate such a host switch. The complexity of the problem suggested that in addition to the singular signatures, there might be a combinatorial use of such genomic features, in nature, defining adaptation to hosts..

**Results:** We used computational rule-based modeling to identify combinatorial sets of interacting amino acid (aa) residues in 12 proteins of IAVs of H1N1 and H3N2 subtypes. We built highly accurate rule-based models for each protein that could differentiate between viral aa sequences coming from avian and human hosts,. We found 68 combinations of aa residues associated to host adaptation (HAd) on HA, M1, M2, NP, NS1, NEP, PA, PA-X, PB1 and PB2 proteins of the H1N1 subtype and 24 on M1, M2, NEP, PB1 and PB2 proteins of the H3N2 subtypes. In addition to these combinations, we found 132 novel singular aa signatures distributed among all proteins, including the newly discovered PA-X protein, of both subtypes. We showed that HA, NA, NP, NS1, NEP, PA-X and PA proteins of the H1N1 subtype carry H1N1-specific and HA, NA, PA-X, PA, PB1-F2 and PB1 of the H3N2 subtype carry H3N2-specific HAd signatures. M1, M2, PB1-F2, PB1 and PB2 of H1N1 subtype, in addition to H1N1 signatures, also carry H3N2 signatures. Similarly M1, M2, NP, NS1, NEP and PB2 of H3N2 subtype were shown to carry both H3N2 and H1N1 HAd signatures.

**Conclusions:** To sum it up, we computationally constructed simple IF-THEN rule-based models that could distinguish between aa sequences of virus particles originating from avian and human hosts. From the rules we identified combinations of aa residues as signatures facilitating the adaptation to specific hosts. The identification of combinatorial aa signatures suggests that the process of adaptation of IAVs to a new host is more complex than previously suggested. The present study provides a basis for further detailed studies with the aim to elucidate the molecular mechanisms providing the foundation for the adaptation process.

## Background

IAVs have been known for a long time to cause disease in a wide range of host species, including humans and various animals. The IAVs are zoonotic pathogens that can infect a broad range of animals from birds to pigs and humans. The interspecies transmission requires that IAVs adapt to the new host and the whole process is facilitated by their high mutation rates [1]. This can result in epidemics and pandemics with severe consequences for both human and animal life. In addition to the yearly epidemics that has proved fatal for at least 250,000 humans worldwide, in the 20^th^ century alone [2], there has been at least five major pandemics; the Spanish flu of 1918, Asian influenza of 1957, Hong Kong influenza of 1968, the age restricted milder Russian flu of the 1977 [3, 4] and the Swine flu of 2009. Thus, new flu epidemics and pandemics are a constant threat. Given our poor understanding of the HAd process of the virus, which can be a major factor for such epidemics and pandemics, it is very hard to predict the type of the virus that will cause the coming outbreaks.

The IAVs are usually classified into subgroups based on the two surface glycol-proteins, hemagglutinin (HA) and neuraminidase (NA). To date, 18 types of HA (H1-H18) and 11 types of NA (N1-N11) are known [5-7]. Most of these species have wild birds as their natural hosts. IAVs are usually adapted and relatively restricted to a single host but occasionally the virus can jump and adapt to a new host species. This cross of the species barrier is proved by the pandemic H1N1, H3N2, H2N2 and the most recent H5N1 and H7N9 subtype outbreaks, which are thought to have evolved from avian or porcine sources [8, 9, 5].

The HA protein plays a crucial part in defining the adaptation of the virus to different hosts since it binds to the receptor providing the entry into host cells. The avian strains of the IAVs are known to prefer a receptor with α2,3-sialic acid linkages while the human strains have a preference for a receptor with α2,6-sialic acid linkages [10]. However, other proteins such as the polymerase subunits have also previously been shown to play a role in the adaptation of IAVs to different hosts [11, 12]. Computational methods, like artificial neural networks, support vector machines and random forests, have been used previously to predict hosts of IAVs [13-15]. Furthermore, several other studies have previously been carried out predicting genomic signatures specifying different hosts, both computationally and experimentally [16-22]. Amino acid changes taken one at a time, i.e. singular aa changes), in viral protein sequences between different hosts have been reported by these studies as host-specific signatures, either directly or indirectly facilitating the HAd process. Despite these findings, this process of adaptation of IAVs in different hosts is still not completely understood. Given the complex nature of the problem we suspected that the HAd signatures are not necessarily univariate. Essentially, in addition to the proven effects of singular aa residues, there might be a combinatorial use of aa residues in nature that affect the adaptation of IAVs to new hosts. To this end, for both H1N1 and H3N2 subtypes, we analyzed aa sequences of 12 proteins expressed by the viruses. We built high quality rule-based models, based on rough sets [23], for each of the 12 proteins, predicting hosts from protein sequences. The models consisted of simple IF-THEN rules that lend themselves to easy interpretation. The combinations of aa residues used by the rules were identified as HAd signatures. In additions to such combinatorial signatures, novel singular signatures were also identified from the rules. The singular and, especially, the combinatorial signatures provide novel insights into the complex HAd process of the IAVs.

## Results

### Feature selection reduces the number of features needed to discern between hosts

Monte Carlo Feature Selection (MCFS) [24]was used to obtain a ranked list of significant features, here significantly informative aa positions in all the proteins for both subtypes, that best discern between the hosts. This step helped us remove any kind of noise that could have been in the data. More importantly, the use of MCFS considerably reduced the number of aa positions to be analyzed further, as shown in Table 1. The HA protein had 628 positions to start with and after running MCFS on the data, we were left with 115 and 88 positions for H1N1 and H3N2 subtypes, respectively (81.7% and 86% reduction in the number aa positions). On average there was a 79.8% reduction in the number of aa positions across all the proteins for H1N1 subtype and 82.8% for the H3N2 subtype (Table 1). Only the significant features were used for further analysis in this study. The ranked lists of the significant features are provided as a supplementary file (see Additional file 1).

**Table 1.**
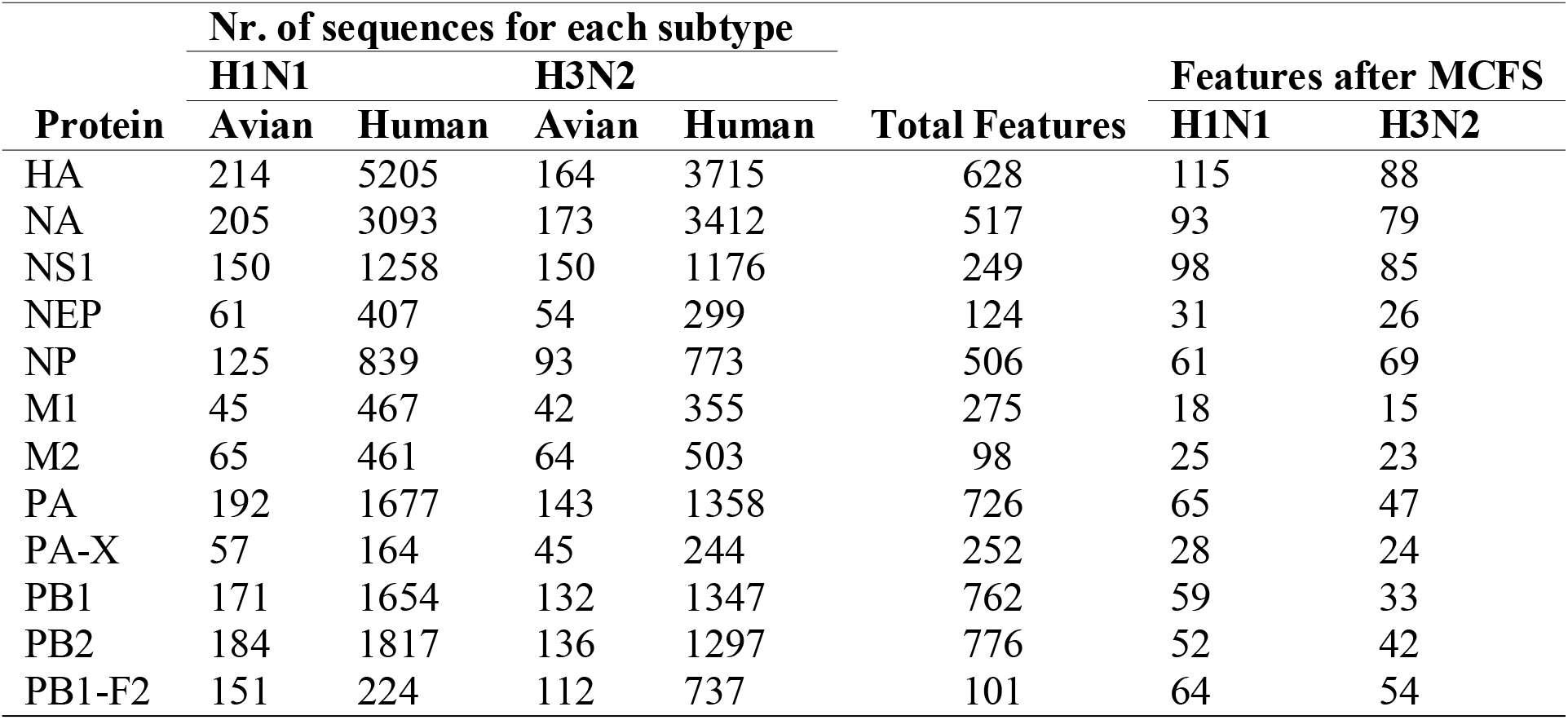
The training data.

### Rule-based models for each protein

Since the number of sequences belonging to human and avian hosts were not balanced in the training data of either subtype (Table 1), we balanced the data sets by a method called under-sampling, as described in detail in Methods. For data sets of each protein and each subtype we created 100 under-sampled subsets. Each of these subsets was used to build a classifier, consisting of IF-THEN rules, whose performance was assessed by a 10-fold cross-validation. Mean accuracies of the 100 classifiers were averaged and shown in Figure 1. HA classifiers for H1N1 and non-structural protein 1 (NS1) classifiers for H3N2 subtypes were the best ones with a mean accuracy of 98% and 98.9%, respectively. Nuclear export protein (NEP) classifiers of the H1N1 subtype and matrix protein 1 (M1) classifiers of the H3N2 subtype had lowest mean accuracy of 83.4% and 88.8%, respectively, which is still a very good result. For each protein of each subtype a single rule-based model containing only the most significant rules from their respective 100 classifiers was inferred (Methods). We then reclassified the training data of each protein with its respective rule-based model to get an idea of its performance in terms of classification of human and avian sequences. Polymerase acidic protein X (PA-X), which is a frame-shift product of the third RNA segment, HA and NEP (NS2) models performed the best (Mathew’s correlation coefficient (MCC) = 1, MCC = 0.99, MCC = 0.99, respectively) among the H3N2 models while HA, NA and NS1 models performed the best among the H1N1 models (MCC = 0.96, MCC = 0.95, MCC = 0.95, respectively) (Figure 2). The poorest of the H1N1 models was the PA-X protein model (MCC = 0.86) and of the H3N2 models was the polymerase basic protein F2 (PB1-F2) protein model (MCC = 0.86). The complete HA H1N1 rule-based model is shown in Table 2. Models for the remaining proteins for both subtypes are provided as supplementary material (Additional file 2).

**Figure 1.**
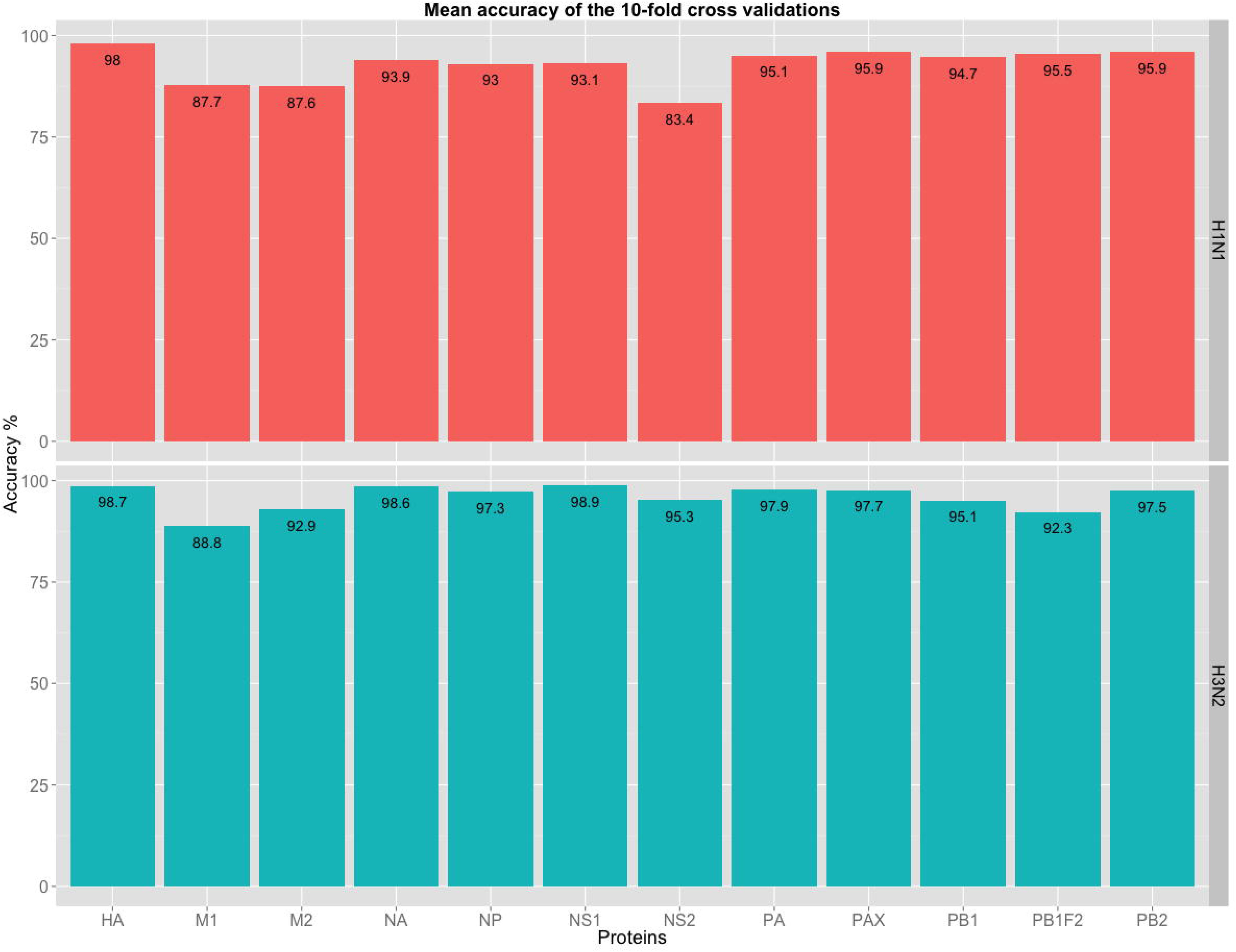
Mean accuracies of the classifiers from 10-fold cross validations. The red bars are for the H1N1 subtype and cyan bars are for the H3N2 subtype.

**Figure 2.**
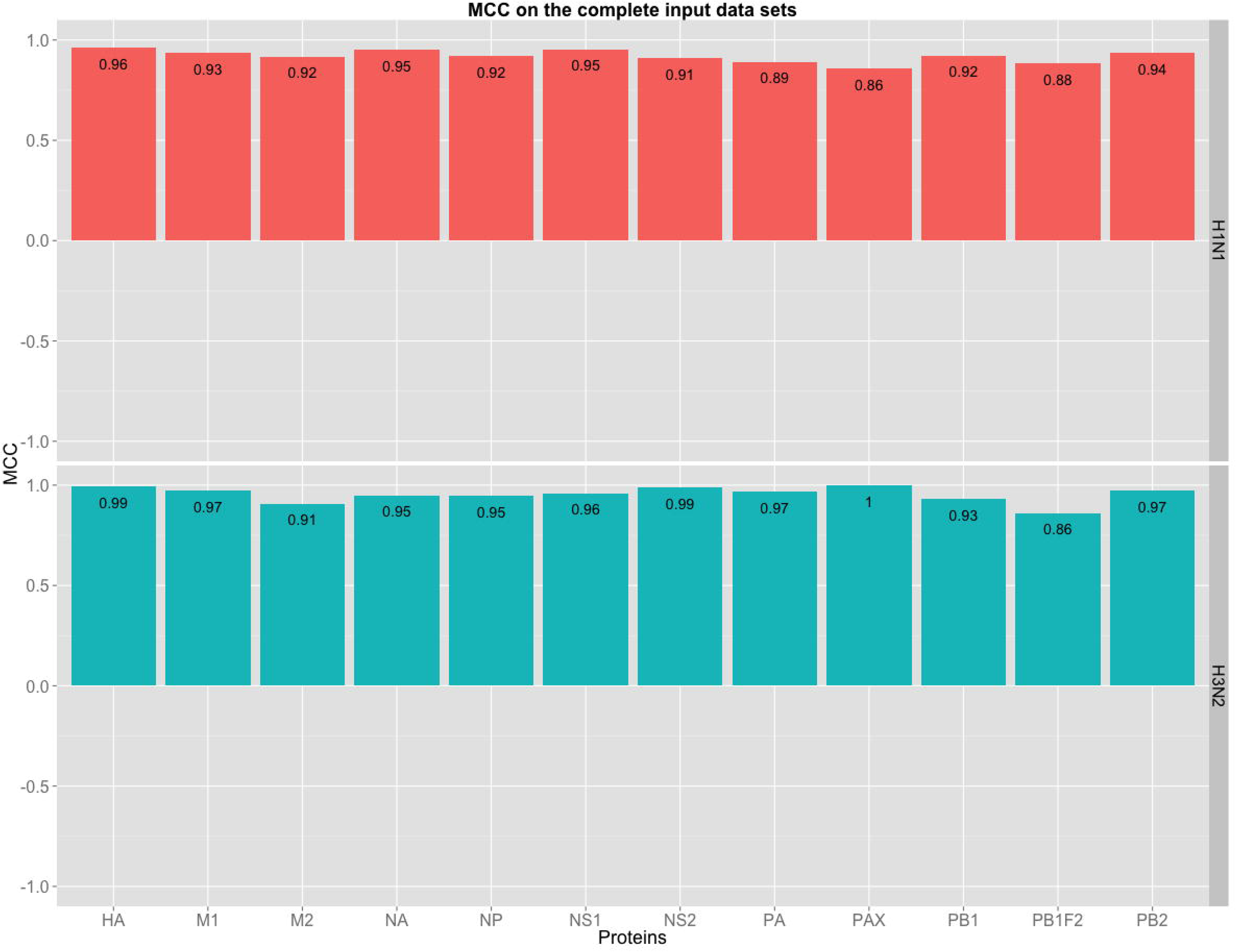
Performance of the rule-based models. The figure shows how well the models perform from a classification point of view, which is shown in terms of Mathew’s correlation coefficient (MCC) values when tested on its corresponding complete input data set for each protein model of both subtypes. A value of 1 means a perfect classification, 0 is for a prediction no better than random and −1 indicates a total disagreement between predictions and observations. The red bars are for the H1N1 subtype and cyan bars are for the H3N2 subtype.

**Table 2.** Rule-based model of HA protein for the H1N1 subtype

| Rule | Accuracy (%) | Support | Decision Coverage (%) |
| --- | --- | --- | --- |
| IF P435=I THEN host=Human | 99.9 | 5128 | 98.4 |
| IF P200=S THEN host=Human | 99.9 | 4052 | 77.8 |
| IF P10=Y THEN host=Human | 99.8 | 3998 | 76.7 |
| IF P88=S THEN host=Human | 99.9 | 3989 | 76.5 |
| IF P6=V THEN host=Human | 99.8 | 3936 | 75.5 |
| IF P222=R THEN host=Human | 99.9 | 3823 | 73.4 |
| IF P220=T THEN host=Human | 100.0 | 3584 | 68.8 |
| IF P516=K THEN host=Human | 99.9 | 1818 | 34.9 |
| IF P200=P and P222=K THEN host=Avian | 91.3 | 229 | 97.7 |
| IF P130=K THEN host=Avian | 91.3 | 218 | 93.0 |
| IF P2=E and P222=K THEN host=Avian | 96.2 | 208 | 93.5 |
| IF P137=A and P544=L THEN host=Avian | 96.1 | 205 | 92.1 |
| IF P78=L and P435=V THEN host=Avian | 97.1 | 204 | 92.5 |
| IF P9=F THEN host=Avian | 98.5 | 204 | 93.9 |
| IF P6=F THEN host=Avian | 98.2 | 169 | 77.6 |
| IF P14=V THEN host=Avian | 99.4 | 165 | 76.6 |
| IF P173=T THEN host=Avian | 98.7 | 158 | 72.9 |

To further verify the validity of the rule-based models created, we tested them on new, unseen data. This data was protein sequences published at the NCBI resource between 30^th^ of November 2014 and 16^th^ of April 2015. For the H1N1 subtype, the rule-based models of M1, nucleoprotein (NP), NS1, NEP (also called non-structural protein 2 (NS2)), PB1-F2, polymerase basic protein 1 (PB1) and polymerase basic protein 2 (PB2) provided perfect classification (i.e. all the sequences were correctly classified). For the H3N2 subtype data, the models of HA, M1, NP, NS1, NEP (NS2), polymerase acidic protein (PA), PB1 and PB2 also gave a perfect classification. Table 3 shows the performance of all rule-based models on the unseen data. A list of names of the viruses that could not be classified or were miss-classified for both subtypes is given in Additional file 3.

**Table 3.** Performance of the rule-based models on the new, unseen data

| Protein | Human Sequences |  | Avian Sequences |  |
| --- | --- | --- | --- | --- |
|  | Total | Correctly classified | Total | Correctly classified |
| HA-H1N1 | 107 | 104 | 2 | 2 |
| HA-H3N2 | 72 | 72 | 4 | 4 |
| M1-H1N1 | 24 | 24 | 0 | 0 |
| M1-H3N2 | 7 | 7 | 0 | 0 |
| M2-H1N1 | 30 | 26 | 1 | 1 |
| M2-H3N2 | 21 | 15 | 3 | 3 |
| NA-H1N1 | 32 | 32 | 2 | 1 |
| NA-H3N2 | 45 | 45 | 4 | 3 |
| NP-H1N1 | 13 | 13 | 1 | 1 |
| NP-H3N2 | 7 | 7 | 4 | 4 |
| NS1-H1N1 | 30 | 30 | 2 | 2 |
| NS1-H3N2 | 18 | 18 | 3 | 3 |
| NEP-H1N1 | 11 | 11 | 2 | 2 |
| NEP-H3N2 | 7 | 7 | 2 | 2 |
| PAX-H1N1 | 17 | 13 | 2 | 2 |
| PAX-H3N2 | 6 | 6 | 0 | 0 |
| PA-H1N1 | 33 | 28 | 2 | 2 |
| PA-H3N2 | 23 | 23 | 3 | 3 |
| PB1F2-H1N1 | 2 | 2 | 2 | 2 |
| PB1F2-H3N2 | 8 | 7 | 4 | 0 |
| PB1-H1N1 | 27 | 27 | 0 | 0 |
| PB1-H3N2 | 19 | 19 | 1 | 1 |
| PB2-H1N1 | 28 | 28 | 2 | 2 |
| PB2-H3N2 | 15 | 15 | 3 | 3 |

### Predicted signatures of HAd

The rule-based models allowed us to further interpret them and see how they differentiated viral avian from viral human sequences. Each of the models was analyzed separately for HAd signatures. The constituent rules of a model associated aa residues at specific positions with an avian or human host. The confidence in these associations is shown as the accuracy, support and the decision coverage shown in the rule-based models. For the combinations in our models we also calculated a combinatorial accuracy gain (CAG), which is the percentage points gain in accuracy of the combination as compared to the average of the accuracies of its constituent singular conditions when taken independently.

### Combinatorial signatures

As expected we found aa combinations in HA, M1, matrix protein 2 (M2), NP, NS1, NEP (NS2), PA, PA-X, PB1 and PB2 proteins to be associated with specific hosts in the H1N1 subtype. In the H3N2 subtype, we found combinations in M1, M2, NEP, PB1 and PB2 proteins. A complete set of combinations for both subtypes is given in a supplementary file (see Additional file 4: Combinations_from_rules). Ciruvis diagrams [25] for visualization of combinations of interacting amino acids were used to illustrate the cases of three or more combinations in the models of both subtypes associated with the avian hosts (see Figure 3 and Figure 4).

**Figure 3.**
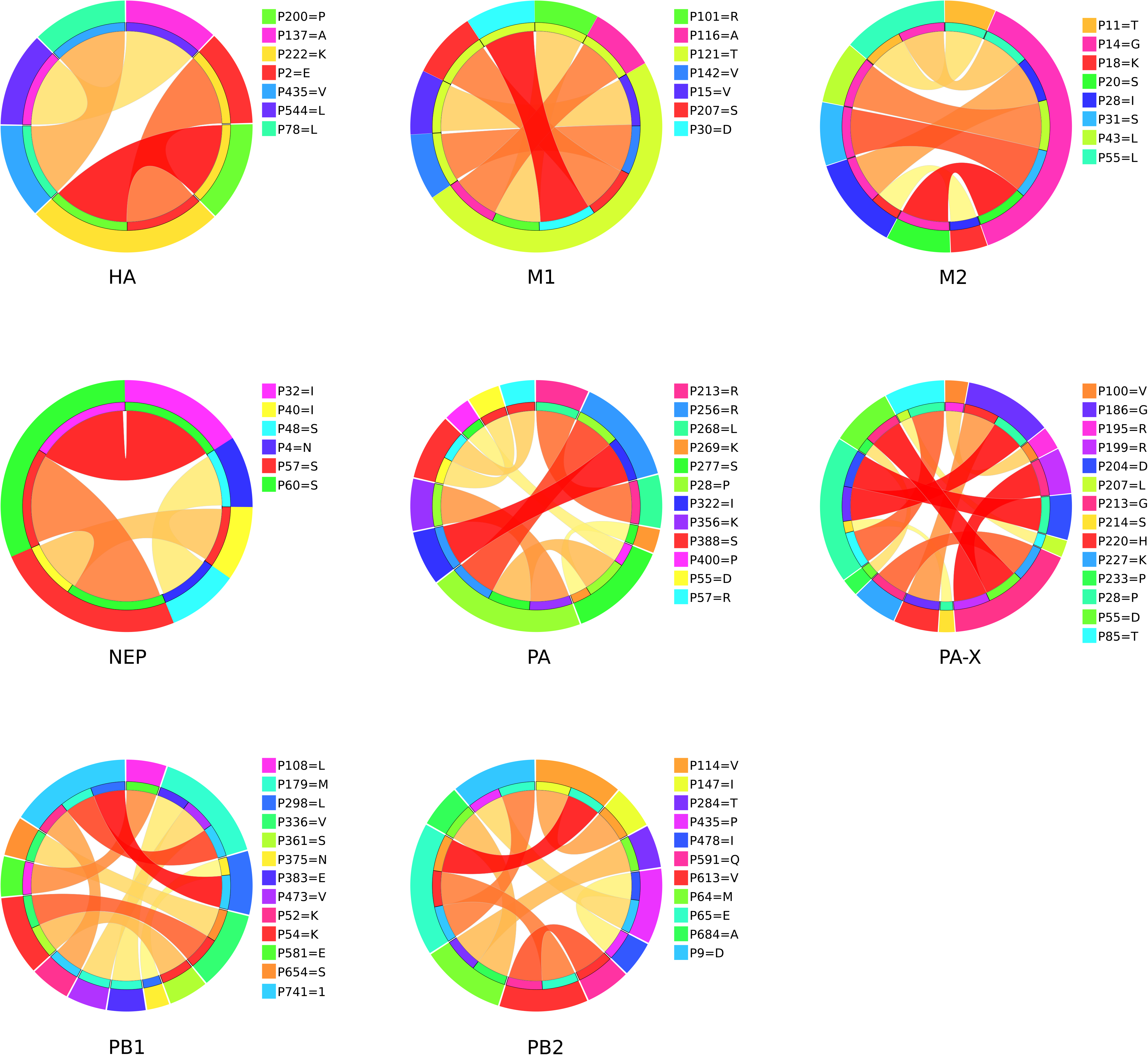
Ciruvis diagrams of combinations from the rules of H1N1 models. Models having at least three combinations are shown. The outer circle shows the positions. The inner circle shows the position or positions to which the position of the outer circle is connected. The edges show these connections. The width and color of the edges are related to the connection score (low = yellow and thin, high = red and thick). The width of an outer position is the sum of all connections to it, scaled so that all positions together cover the whole circle [25].

**Figure 4.**
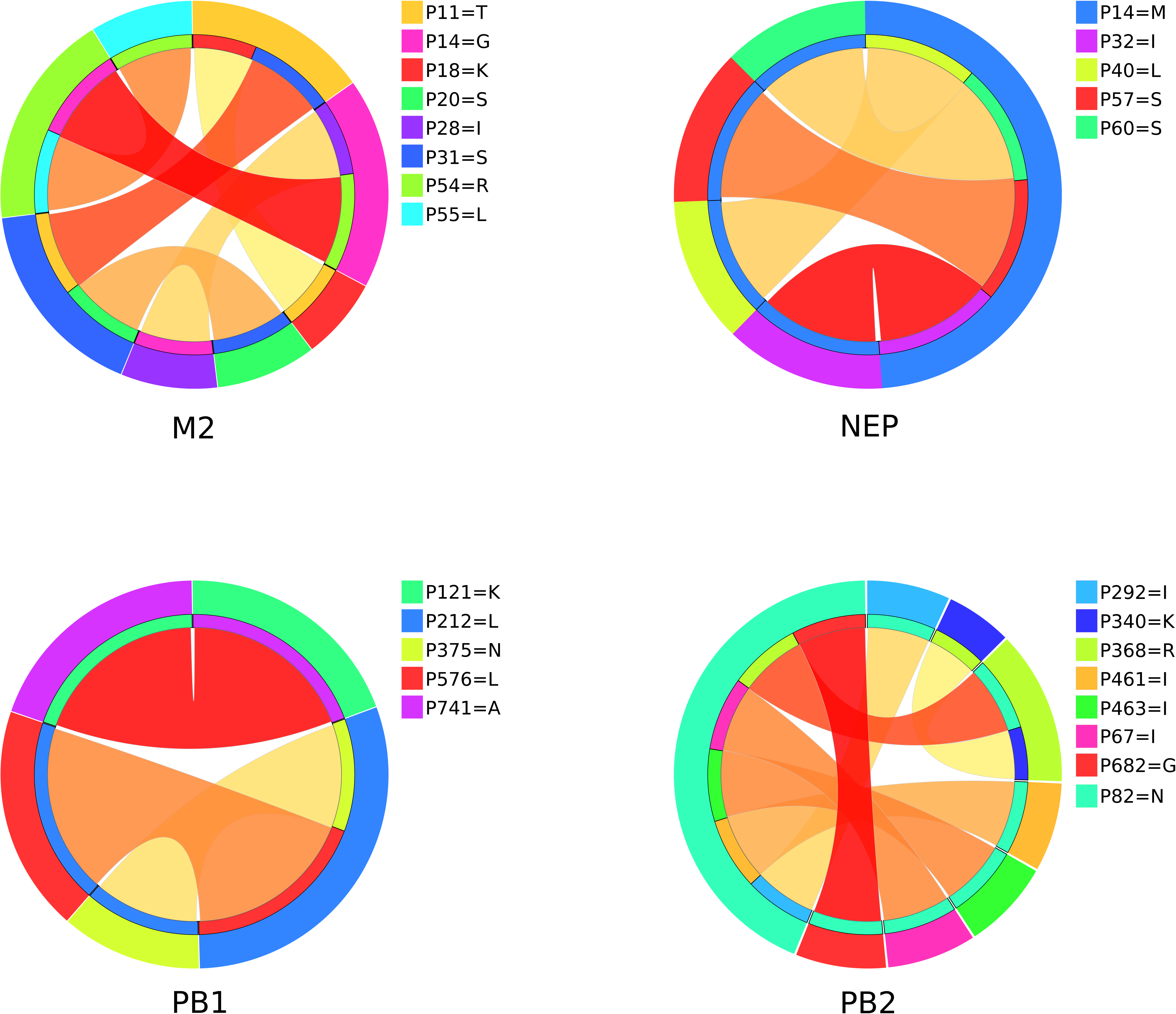
Ciruvis diagrams of combinations from the rules of H3N2 models. Models having at least three combinations are shown. The outer circle shows the positions. The inner circle shows the position or positions to which the position of the outer circle is connected. The edges show these connections. The width and color of the edges are related to the connection score (low = yellow and thin, high = red and thick). The width of an outer position is the sum of all connections to it, scaled so that all positions together cover the whole circle [25].

Residues 14G of the M2 H1N1 model and 82N of the PB2 H3N2 model were the most connected ones interacting with six other aa residues each. Amino acid residues having interactions with more than one other residue, in both the subtypes are listed in Table 4. These strongly interacting residues might be relatively more essential to HAd than the less connected ones.

**Table 4.** Amino acid residues having the most interactions in the models of both subtypes.

| Subtype | Protein | Positions | Number of interactions |
| --- | --- | --- | --- |
| H1N1 | HA | 222K | 2 |
|  | M1 | 121T | 5 |
|  | M2 | 14G | 6 |
|  | NEP | 57S, 60S | 2 |
|  | PA | 28P, 277S | 3 |
|  | PA-X | 28P | 4 |
|  | PB1 | 179M, 741A | 3 |
|  | PB2 | 65E | 3 |
| H3N2 | M1 | 101R | 2 |
|  | M2 | 11T, 14G, 31S, 54R | 2 |
|  | NEP | 14M | 4 |
|  | PB1 | 212L | 2 |
|  | PB2 | 82N | 6 |

### Singular (linear) signatures

Previous studies [16-22] mostly found the adaptation signatures on the internal proteins and did not look into surface glycoproteins (HA and NA). In contrast, we found singular signatures on all the proteins of both subtypes, including the HA, NA and the newly discovered PA-X proteins. PA-X protein shares the human signature 85I with PA in the H1N1 model while it shares human signatures 28L and avian signature 28P in the H3N2 models. In total, 189 singular signatures were found, in both subtypes combined. Out of these, 132 signatures were novel and not reported by the previous studies (Table 5). A complete list of singular signatures is given in the supplementary material (see Additional file 4: singletons_H3N2, singletons_H1N1)

**Table 5.** Novel singular aa positions associated to host adaptation

| Protein | Novel singular positions |
| --- | --- |
| HA | 6,9,10,14,23,47,66,69,78,88,91,94,130,173,189,200,220,222,435,516 |
| M1 | 30,116,142,207,209 |
| M2 | 13,16,31,36,43,51,54 |
| NA | 16,18,19,23,30,40,42,44,46,47,74,79,147,150,157,166,232,285,341,344,351,369,372,389,397,435,437,466 |
| NP | 31,53,98,146,444,450,498 |
| NS1 | 6,7,14,23,27,28,74,123,152,192,220,226 |
| NS2 | 6,7,14,32,34,48,83,86 |
| PA | 85,323,336,348,362,300 |
| PAX | 28,85,210,233 |
| PB1 | 12,54,59,113,175,212,339,435,576,586,587,619,709 |
| PB1F2 | 3,6,12,17,21,25,26,27,28,33,47,52,54,57,58,60,62,65,82 |
| PB2 | 54,65,354 |

### Specific aa changes associated with HAd

Some of the rules from our models associated different residues at the same aa positions with avian and human hosts. This can be seen as a mutation (aa change) associated to the adaptation of the viral proteins to a specific host. Eight mutations were found for the H1N1 subtype and 10 for the H3N2 one. In the H1N1 subtype, mutations F6V in HA, P46T and L74V in NA, I6M in both NS1 and NEP and L58- in PB1-F2 were novel. In the H3N2 subtype, mutations R78E in HA, A30I, N40Y and I44S in NA, P28L and R57Q in PA and P28L in PA-X were not identified in the previous studies. Table 6 shows all such mutations in both subtypes.

**Table 6.** Amino acid changes associated with host adaptation

| H1N1 |  |  |  | H3N2 |  |  |  |
| --- | --- | --- | --- | --- | --- | --- | --- |
| Protein | Position | Avian | Human | Protein | Position | Avian | Human |
| HA | 6 | F | V | HA | 78 | R | E |
| NA | 46 | P | T | NA | 30 | A | I |
|  | 74 | L | V |  | 40 | N | Y |
| NP | 100 | R | I,V |  | 44 | I | S |
| NS1 | 6 | I | M | NP | 16 | G | D |
| NEP | 6 | I | M | PA-X | 28 | P | L |
| PB1-F2 | 58 | L | - | PA | 28 | P | L |
| PB2 | 588 | A | I |  | 57 | R | Q |
|  |  |  |  | PB2 | 9 | D | N |
|  |  |  |  |  | 64 | M | T |

### Predicted signatures are not specific to sub-clades of the strains

The support and the decision coverage of the rules showed whether the aa signatures identified were specific to sub-clades or were more general i.e. spread out across the sub-clades. The higher decision coverage indicated more generality of the rule. For example, the top five rules for the avian class have the following very high decision coverage: rule1 – 98.5%, rule2 – 98.5%, rule3 – 97.8%, rule4 – 98.5% and rule5 – 97.8%. It follows that the rules are general. To further illustrate this generality, and to show the diversity in our training data set, a phylogenetic analysis was carried out (additional file 5). Top five rules specifying each host were mapped onto the created phylogenetic trees, separately for each host, for all the proteins of both subtypes. As an example, consider the avian PB2 H3N2 tree (Figure 5). 91.4% of the sequences are covered by rule 1, 2, 3, 4 and 5, which is illustrated by the violet coloring of the leaves in the tree. Only, 1.4% of the sequences are not covered by rule4, yet they are covered by rule 1, 2, 3, and 5, and similarly for the remaining coverage. For the corresponding human tree, the figures are 89.3% coverage for the top five human rules. One can see that this generality prevails in all other proteins.

**Figure 5.**
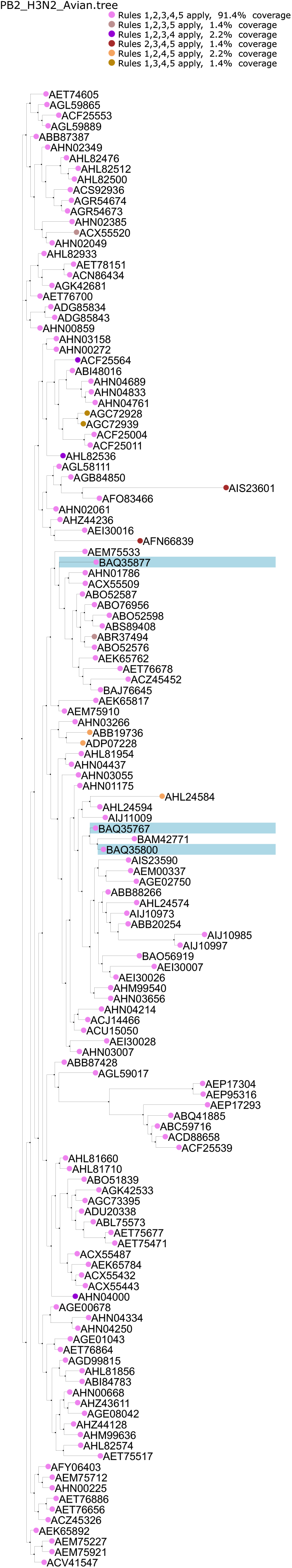
Phylogeny of PB2 H3N2 protein of avian hosts annotated with top 5 avian rules form the PB2 H3N2 model. Each sequences is represented by its GeneBank accession. The violet nodes mark the sequences that supports rule 1,2,3,4 and 5, which are 91.4% of the total sequences. Similarly the DarkViolet nodes mark the sequences that support rule 1, 2, 3 and 4 but lacks support for rule 5, which are 2.2% of the total sequences. The nodes with a LightBlue background are the new, unseen sequences. The unmarked nodes do not support the top 5 rules, and were either supporting rules other than the top 5 or were not classified by the models.

### Validity of HAd signatures across H1N1 and H3N2 subtypes

To see whether the signatures associated with HAd identified in the H1N1 subtype could also function as signatures for the H3N2 subtype and vice versa, we classified H3N2 subtype data with H1N1 models and H1N1 subtype data with H3N2 models. Good classifications meant that the rules (and consequently the signatures associated to adaptation) generated for one subtype were valid for the other one. Bad classifications meant that the rules of one subtype did not hold for the data of the other subtype and hence no cross-subtype marker validity. Both HA and NA H1N1 models were bad classifiers for the HA and NA of the H3N2 type data, respectively since they failed to distinguish avian sequences in the data in both cases (Sp = 0) (Table 7). It should be kept in mind that the outcome *human* was considered positive outcome and the outcome *avian* considered as a negative one. The PA-X H1N1 model could not recognize human sequences in the PA-X H3N2 data (Sn = 0). Furthermore, the models of PA, PB1-F2 and PB1 proteins of H1N1 subtype were bad classifiers of the H3N2 data (MCC = −0.11, MCC = 0.056, MCC = 0.302), specifically failing to identify sequences coming from human hosts (Sn = 0.021, Sn = 0.023, Sn = 0.563). This meant that H1N1 HAd signatures in the models of HA, NA, PA-X, PA, PB1-F2 and PB1 proteins were not valid for H3N2 subtype data and these proteins of the H3N2 subtype carried only H3N2-specific HAd signatures. Contrary to this, the H1N1 models of M1, M2, NP, NS1, NEP and PB2 proteins were able to distinguish between H3N2 subtype sequences coming from avian and human sources reasonably well (Sn = 0.97–1.0; Sp = 0.64–0.94; MCC = 0.776–0.941). It proved that these proteins of the H3N2 subtype, in addition to the stronger H3N2 HAd signatures, also carried H1N1 HAd signatures.

**Table 7.** Performance of the H1N1 models on H3N2 data and vice versa. Sensitivity is the ability to correctly predict human sequences and specificity is the ability to correctly predict avian sequences where 1 means perfect prediction and 0 means no correct predictions. Mathew’s correlation coefficient (MCC) value is a measure of how well the model performs overall where 1 is perfect prediction, 0 is similar to prediction by chance and −1 is total disagreement between observations and predictions. “na” means the measure could not be calculated for the given model.

|  | Protein | Sensitivity | Specificity | MCC |
| --- | --- | --- | --- | --- |
| **H3N2 data - H1N1 models** | HA | 1 | 0 | na |
|  | M1 | 1 | 0.895 | 0.941 |
|  | M2 | 1 | 0.742 | 0.848 |
|  | NA | 1 | 0 | na |
|  | NP | 1 | 0.891 | 0.938 |
|  | NS1 | 1 | 0.745 | 0.849 |
|  | NEP | 1 | 0.642 | 0.776 |
|  | PA-X | 0 | 1 | na |
|  | PA | 0.021 | 0.93 | -0.11 |
|  | PB1-F2 | 0.023 | 1 | 0.056 |
|  | PB1 | 0.563 | 0.909 | 0.302 |
|  | PB2 | 0.979 | 0.949 | 0.873 |
| **H1N1 data - H3N2 models** | HA | 0 | na | na |
|  | M1 | 0.957 | 0.975 | 0.885 |
|  | M2 | 0.987 | 0.766 | 0.804 |
|  | NA | 1 | 0 | -0.004 |
|  | NP | 0.363 | 0.984 | 0.251 |
|  | NS1 | 0.365 | 0.993 | 0.236 |
|  | NEP | 0.027 | 1 | 0.06 |
|  | PA-X | 0.202 | 0.982 | 0.224 |
|  | PA | 0.247 | 0.995 | 0.177 |
|  | PB1-F2 | 0.991 | 0.804 | 0.831 |
|  | PB1 | 0.992 | 0.877 | 0.888 |
|  | PB2 | 0.956 | 0.951 | 0.788 |

The H3N2 models of HA, NA, NP, NS1, NEP, PA-X and PA proteins could not classify avian and human sequences of H1N1 subtype correctly (MCC = −0.004– 0.251). This means that these proteins of the H1N1 subtype carried H1N1-specific signatures. Whereas the successful classifications of H1N1 subtype data of M1, M2, PB1-F2, PB1 and PB2 proteins by the respective H3N2 models (MCC = 0.788–0.888; Sn = 0.956–0.992; Sp = 0.766–0.951) proved that these H1N1 proteins carried both H1N1 and H3N2 signatures.

## Discussion

In this study we have focused on H1N1 and H3N2 and restricted our analyses to these two subtypes. Our models performed reasonably well since all of them had an average accuracy of more than 90% in the 10-fold cross validation except NEP (NS2), M1 and M2 protein models of the H1N1 type (Accuracy: 83.4%, 87.7% and 87.6%, respectively) and M1 protein model of the H3N2 type (Accuracy 88.8%) (Figure 1). The reason for the relatively low accuracies of the above exceptions could be either the lack of training sequences from which the models learn or these sequences may lack stronger genomic signatures specific to hosts.

In previous studies [16-22], signatures of adaptation were mostly found on the internal proteins, especially in viral ribonucleoprotein complexes consisting of viral polymerases and NP. The fact that we were able to build high quality models for all the proteins for both subtypes, indicated that all the proteins, including the highly variable HA and NA proteins and the recently discovered PA-X protein, carry genomic signatures specific to hosts. A major difference between our models and the ones previously reported is that the previous models were black box classifiers whereas our models are transparent. Black box classifiers give classification but do not provide any straightforward possibility to identify which parameters and for which values a classification is obtained. Transparent classifiers allow explicit analysis of the model, i.e. the features and their values, for each classified object. The models created in this study used aa positions as features and aa residues at those positions as the values for those features, hence lending themselves for easy interpretation and further analysis.

Previous studies listed above reported only on singular aa positions as HAd signatures. However, in addition to singular aa positions, we also identified combinations of aa residues at specific positions as HAd signatures. This is the very first time that combinations of aa positions are reported in this context. These combinations are shown as conjunctive rules, i.e., rules with more than one condition in the IF part. It appeared that some aa residues were part of more than one combination in our models. This may suggest that these residues are relatively more important in establishing HAd then the ones appearing in one combination only (Table 4).

In the M2 H1N1 model, the combinations associated with avian hosts had a Glycine (G) residue at position 14 while the combinations for human hosts had a Glutamic acid (E) in the same position. Similarly, in PB2 H3N2 model, Arginine (R) at position 340 was associated to avian hosts while Lysine (K) residue at the same position to human hosts. It seems that the mutations G14E in M2 H1N1 and R340K in PB2 H3N2 model facilitate the shift of hosts from avian to human. However, these residues always appear in combination with other residues and therefore they cannot be used in forms other than the combinations themselves. The reason is obvious. The confidence measures (accuracy, support and decision-coverage) were calculated for the combination as a whole. We do not report such mutations in our list of mutations affecting HAd although they indicate an effect. The functions of these combinations at a molecular level are not understood yet, but they provide a novel and interesting perspective of looking at sequence based HAd signatures.

HA and NA of both subtypes were found to be only carrying subtype-specific signatures. This goes well with the current knowledge that these two proteins are the most diverse proteins that are specifically adapted to interact with the host cell. M1, M2 and PB2 are shown to be the most conserved proteins from the point of view of host specifying genomic signatures since they carried the host signatures valid for both subtypes.

The signatures found in this study were also considered in other contexts in other studies such as viral viability and antiviral resistances. For instance, positions 30, 142, 207 and 209 occurring in the H1N1 M1 models have been previously shown to affect viral production when mutated [26], while mutation S31N derived from M2 models is a known marker of amantadine resistance [27-30]. Table 8 lists all the aa residues and their descriptions as found in different contexts in the literature. All these different contexts, that the aa residues from our models are described in, show that they affect the fitness of the viruses in one or the other way, which in turn facilitates their adaptation to the new environment or hosts.

**Table 8.** Amino acid positions discussed in literature from the models of both the subtypes for all proteins

| Protein | Positions | Description |
| --- | --- | --- |
| M1 | 115,121,137 | Known signatures of host-adaptation [18, 21, 22] |
|  | 30,142,207,209 | Affecting viral production on mutation [26] |
|  | 121 | Affecting viral replication [44] |
|  | 101 | Determinant of temperature sensitivity [45], located in a transcription inhibition site [46] and is also interacting with NEP [47] |
| M2 | 11,14,18,20,28,55,57,78,82,89,93 | Known signatures of host-adaptation [18, 21, 22, 48] |
|  | 31 | S31N is a known marker for amantadine resistance [27-30] |
|  | 18,20 | Lie next to 17,19 which forms a di-sulphide bond [49] |
| NS1 | 18,21,22,53,60,70,81,112,114,171,215,227 | Known signatures of host-adaptation [19, 50, 17, 21, 22, 20] |
|  | 215 | Required for Crk/CrL-SH3 binding [51] |
|  | 123 | Necessary for interaction with PKR, resulting in an inhibition of eIF2alpha phosphorylation [52] |
|  | 95 | Along with others, has been shown to be necessary for binding p85beta and activating PI3K signaling [53, 54] |
|  | 220 | Part of nuclear localization signal 2 essential for the importin-alpha binding [55] |
| NEP(NS2) | 57,60,70,107 | Known signatures of host-adaptation [17, 56, 18, 22, 21] |
| NP | 16,33,100,214,283,313,351,353,357,422 | Known signatures of host-adaptation [20, 18, 57, 19, 22, 21] |
|  | 16 | D16G shown to decrease pathogenicity several fold [58] |
| PA | 28,55,57,65,256,268,277,356,382,400,409 | Known signatures of host-adaptation [19, 18, 20, 57, 22, 21] |
|  | 85,336 | Residues 85I and 336M are deemed important for enhanced polymerase activity in mammalian cells [59] |
|  | 57,65,85 | Shown to be involved in suppressing the host cell protein synthesis during infection [60] |
| PB1 | 52,179,216,298,327,336,361,375,581,741 | Known signatures of host-adaptation [57, 18, 21, 22, 16] |
|  | 581 | Shown to be conferring temperature sensitivity to human influenza virus vaccine strains [61] |
|  | 473 | Mutation at position 473 has been shown to decrease polymerase activity [62] |
| PB2 | 9,44,64,81,105,271,292,368,453,588,613,682,684 | Known signatures of host-adaptation [57, 18, 19, 22, 21] |
|  | 591 | 591Q is known to mimic the effect of 627K [63, 64] |
|  | 271 | 271A shown to increase polymerase activity in mammalian cells [65] |
|  | 271,588 | Also been shown to be host range determinants [66] |
| PB1-F2 | 16,23,42,66,70,73,76 | Known signatures of host-adaptation [17, 22] |
|  | 66 | Linked with affecting pathogenicity [67] |
| NA | 46,47,74,147,15 | Under selection pressure with a shift of hosts from birds |
|  | 7,341,351 | to humans [57] |
|  | 344 | Calcium ion binds here that stabilizes the molecule (UniProt: Q9IGQ6). |
| HA | 2,6,9,10,14 | Signal peptide domain |
|  | 88,173,220,22 | Position 71, 159, 206 and 208 of the fully-mature HA with H3-numbering [68]) are part of the antigenic sites Cb, Sb and Ca of the HA protein, respectively [69, 70] |

## Conclusions

The highly predictive rule-based models built for 12 proteins for H1N1 and H3N2 subtypes suggest that there are HAd signatures on all the protein including the diverse HA, NA and the newly discovered PA-X protein that were not previously studied in this context. In addition, the transparent nature of our method allowed us to further investigate our models for how the predictions are actually done. This resulted in a list of aa residues and their combinations associated with host specifity. Some of the aa residues identified in this study were already known while others are novel. The ability of our methods to capture the combinatorial nature of the HAd process makes this study unique in its nature. We discovered that the surface proteins HA and NA carry subtype-specific host signatures in both subtypes while NP, NS1, NEP, PA-X and PA of the H1N1 subtype and PA-X, PA, PB1-F2 and PB1 of the H3N2 subtype carry subtype-specific host signatures. We showed that M1, M2, PB1-F2, PB1 and PB2 of the H1N1 subtype carried H1N1 and some additional H3N2 signatures, and vice versa, M1, M2, NP, NS1, NEP and PB2 of the H3N2 subtype carried H3N2 and some additional H1N1 host signatures. The computational results presented here will eventually require further analysis by testing the host-pathogen interactions under laboratory conditions. We believe that the computational analyses provide important support in the characterization of host-pathogen interactions and the proper combination of *in silico* and *in vitro* (probably even *in vivo*) studies will yield important novel information concerning the infection biology of various viruses and other infectious agents.

## Methods

The combined feature selection – rule-based modeling methodology used in this is similar to our previous work where we identified a complete map of potential pathogenicity markers in the H5N1 subtype of the avian influenza A viruses [31].

### Data

The data used to make the models was downloaded from the NCBI flu database found at http://www.ncbi.nlm.nih.gov/genomes/FLU/Database/nph-select.cgi?go=database [32]. Full-length plus (nearly complete, may only miss the start and stop codons) protein sequences of the twelve proteins namely, HA, NA, NP, M1, M2, NS1, NEP (NS2), PA, PA-X, PB1, PB2 and PB1-F2, were separately downloaded as published up till November 30, 2014. Identical sequences were represented by the oldest sequence in the database. For each protein, sequences of the H3N2 and H1N1 subtypes of avian and human hosts were downloaded. Sequences of the mixed subtypes were not included in this study. Table 1 shows the number of sequences for each of the proteins for each subtype. For each protein we combined the sequences of the two subtypes used in this study into a single file and aligned them with MUSCLE (v3.8.31) [33].

### Decision Tables

A decision table was created for each of the proteins for both the subtypes. A decision table can be seen as a tabularized form of the aligned FASTA sequences with an extra decision/label column, which in our case was the host information. The first column of the decision tables contained the identifier of the sequence, and the last column was the label/outcome column, the host information in our case and the rest of the columns represented the sequence information corresponding to the aligned FASTA files. The alignment gaps were represented by a ‘?’ in the decision tables. The rows of a decision table were called objects each representing a particular aa sequence and a label. Columns other than the first and the last one were the features.

### Feature selection

MCFS, as described in [24], was used to rank the features of the decision tables with respect to their ability to discern between avian and human hosts. MCFS is implemented as a software package dmLab [34]. MCFS uses a large number of decision trees and assigns a normalized relative importance (RI-norm) score to each feature such that the features contributing more to the discernibility of the outcome gets a higher score. Statistical significance of the RI-norm scores was assessed with a permutation test and significant features (p<0.05), after Bonferroni correction [35], were kept as described in [36]. Only these features were used in the further rule-based model generation.

### Under-sampling the data sets

In the training data for both subtypes, the number of sequences from human hosts was considerably higher than that from the avian hosts. It has previously been shown that this imbalance affects the learning in favor of the dominating class [37]. However to address this problem one can artificially balance the classes [38]. To this end, a technique called under-sampling was used where the sequences belonging to the dominating class were randomly sampled equal to the class having the lesser number of sequences and repeated this step 100 times. In this way for each protein and for each subtype we created 100 subsets where the number of sequences belonging to human and avian hosts were equal. A single rule-based classifier was inferred from each of the subsets, which resulted in 200 classifiers per protein. We illustrate the process with the following example.

The data set of the NA protein of the H1N1 subtype had 3093 human and 205 avian sequences, which was a significant imbalance in the number of sequences. From the human set we created subsets by randomly extracting 100 times 205 human sequences and joining them with the 205 avian sequences to create 100 subsets.

### Rough sets and rule-based model generation

Rough set theory [23] was used to produce minimal sets of features that can discern between the objects belonging to different decision classes. ROSETTA [39], a publicly available software system that implements rough sets theory, was used to transform the minimal sets of features into rule-based models [40] that consisted of simple IF-THEN rules. A complete description of rough sets can be found in [41] and the combined MCFS-ROSETTA approach to model generation in bioinformatics is described in [42].

The input data to ROSETTA were the balanced decision tables created in the previous step with only the significant features obtained from applying MCFS. ROSETTA computed approximately minimal subsets of feature combinations that discerned between avian and human hosts with the Johnsons algorithm implemented in ROSETTA. The classifiers were collections of IF-THEN rules. A sample rule from the HA-H1N1 model: reads as: **“IF** *at position 200 there is a Proline residue* **AND** *at position 222 there is a Lysine residue* **THEN** *the sequence is from an avian host***”**.

| Rule | Accuracy (%) | Support | Decision Coverage(%) |
| --- | --- | --- | --- |
| IF P200=P AND P222=K THEN host=Avian | 91.3 | 229 | 97.7 |

There is additional information about the rules available. *Support* is the set of sequences (229 sequences) that satisfy the conditions of the left hand side (LHS), i.e. the set of sequences that have a proline residue at position 200 and a lysine residue at position 222. For this rule, *Accuracy* is 91.3% that is the proportion of correctly classified sequences to the total number of supporting sequences (209/229). Human sequences are considered positive and avian as negatives in this study. The decision coverage for this rule is 97.7%, which means it correctly classifies 97.7% of the total avian sequences used to train the classifier. It is calculated as follows:

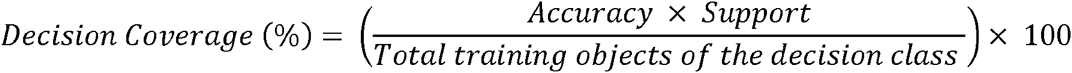

*Accuracy* x *Support* gives us the total number of sequences that are correctly classified by the rule. Since the rule is for the avian decision class, the total number of avian sequences used to train the classifier was 214. So for the stated rule the decision coverage will be ((0.913*229)/214)*100, which is equal to 97.7%. The above rule is a conjunctive rule since there is a conjunction of conditions (`P2 00=P AND P222=K`) in the left hand side (LHS) of the rule. A conjunctive rule captures the underlying combinatorial nature of the HAd process. Each conjunctive rule must always be used as combination only, because the support, accuracy and the decision coverage measures are calculated for the conjunction and not for the individual conjuncts. A rule can also be a singleton rule where LHS consists of only a single condition. The confidence in these classifiers come from the 10-fold cross validation performed in ROSETTA. In a 10-fold cross validation step the input data set is randomly divided into ten equal subsets, say {P1, …, P10}. A classifier is trained on the first nine subsets {P1, …, P9} and then tested on the remaining, P10 subset. In the next run, another classifier is trained on {P1, …, P8, P10} and its performance is tested on the remaining subset, this time P9. Notice that each time the test set is a different one. The process is repeated 10 times and by then each subset has been used once as a test set. The performance of all the classifiers is averaged and presented as a cross-validation accuracy. Such a validation is quite common in machine learning since one becomes more or less assured that the performance of the classifier was not simply by chance.

### Extraction of a single rule-based model for each protein

Rules from all the 100 classifiers were combined into a single file. Duplicates were removed. Among partially identical rules, the one with the highest decision coverage was kept. If the difference of decision coverage was lower than 1% then the shortest (the rule with least conditions) was kept. Accuracy, support and decision coverage were calculated on the complete data set for all the rules. Rules that were below the 90% accuracy and 30% decision coverage thresholds were discarded. In this way we extracted a single, high quality rule-based model for each of the protein for both H1N1 and H3N2 subtype data.

### Classification of sequences

In order to classify a sequence, each rule from the model was applied on it. If the conditions of the rule matched the sequence, the rule was said to fire on the sequence. Every fired rule voted for a particular classification specified by its THEN-part. The number of votes a fired rule casted was the accuracy multiplied by the support of the rule. For a sequence several rules may fire, each casting votes in favor of the class in the THEN-part. The final classification was assigned based on the majority of votes. Consider the rules:

In case of 1) IF P70=S THEN host=Avian. Acc=94.0% Supp=50

2) IF P14=M and P32=I THEN host=Avian. Acc=93.0% Supp=43

3) IF P14=L THEN host=Human. Acc=100% Supp=285

4) IF P57=L THEN host=Human. Acc=100% Supp=273

Now let us assume that these four rules are applied to a sequence an it turns out that Rule 2, 3 and 4 fire for this sequence. Rule 2 will cast 40 (0.93*43) votes for class Avian while rule 2 and rule 3 will cast 285 and 273 votes in favor of class Human. So, the sequence will be classified as class Human since the number of votes is 558 versus 40.

In case of no rules fired or there was a tie in the votes, the sequences were labeled as unknown.

### Performance evaluation statistics of the rule-based models

In this study the outcome *human* was considered as a positive outcome and outcome *avian* was considered as a negative one. True positives (TP) were sequences correctly classified as coming from human hosts. True negatives (TN) were sequences correctly classified as coming from avian hosts. False positives (FP) were actually avian sequences but incorrectly classified as human sequences and false negatives (FN) were actually human sequences that were incorrectly classified as avian sequences. The performance of the models for all the proteins for both H1N1 and H3N2 was assessed by the following statistics.

Sensitivity: it is also known as the true positive rate (TPR). In our case, rate at which a model correctly identifies sequences coming from a human host is the sensitivity i.e. a sequence originally from human host and classified as coming from human hosts by the model. It is calculated with the following formula:

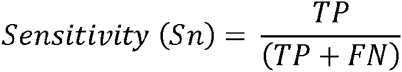

Specificity: Also known as the true negative rate (TNR). The rate at which the model correctly identifies avian sequences is the specificity, which is calculated by:

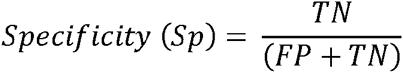

Mathew’s correlation coefficient: It is a measure of how well a model classifies as a whole. The difference with accuracy is that unlike accuracy Mathew’s correlation coefficient is not effected by un-balanced data and hence gives a better overall idea of how well the model is classifying. It is calculated by the following formula:

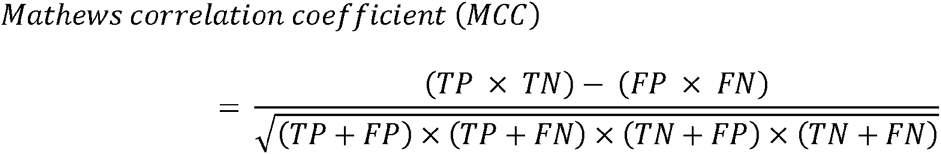

### From alignment positions to true positions

In this study the aa positions for all the H3N2 proteins except the PB1-F2 corresponds to the positions of the *A/<u>Victoria</u>/JY2/1968* virus. For all but PB1-F2 proteins of the H1N1 data, the positions shown in this study correspond to positions on the *A/<u>Wisconsin</u>/301/1976* virus. The PB1-F2 protein for both viruses is in a truncated form and we wanted to show positions from a full-length protein. For this reason we mapped the PB1-F2 H3N2 positions to the PB1-F2 of the *A/New York/674/1995* virus and the PB1-F2 H1N1 positions to full-length PB1-F2 of the *A/duck/<u>Korea</u>/372/2009* virus.

### Phylogenetic analysis

FastTree 2.1.8 [43] was used to create the phylogeny trees.

### Scripting programming language

Python was used for scripting purposes.

## List of abbreviations

aa: Amino acids
CAG: Combinatorial accuracy gain
HA: Hemagglutinin
IAVs: Influenza A viruses
LHS: Left hand side
M1: Matrix protein 1
M2: Matrix protein 2
MCC: Mathew’s correlation coefficient
MCFS: Monte carlo feature selection
NA: Neuraminidase
NEP: Nuclear export protein
NP: Nucleoprotein
NS1: Non structural protein 1
NS2: Non structural protein 2
PA: Polymerase acidic protein
PB1: Polymerase basic protein 1
PB2: Polymerase basic protein 2
Sn: Sensitivity
Sp: Specificity

## Authors Contribution

ZK has performed all computational experiments and together with JK was the main contributor to the paper. MK and SB have contributed the idea to analyze the virus data following the earlier work of JK. They contributed to writing the paper. JK provided the computational methods, supervised the work and together with ZK was the main contributor to the paper.

## Acknowledgements

We would like to thank Husen Umer who provided valuable comments during various stages of the work.

This research was supported by Uppsala University, Sweden, the ESSENCE grant, (ZK and JK), JK was supported in part by Institute of Computer Science, Polish Academy of Sciences, Poland. The EMIDA ERA-NET FP7 EU projects Epi-SEQ (nr. 219235), NADIV (nr. ID 108), the SLU Award of Excellence provided support to SB, and the Swedish Research Council FORMAS Strong Research Environments project, nr 2011-1692, “BioBridges”) to ML and SB.

## Additional Files

**Additional file 1: This file contains the lists of significant features that were selected by MCFS for all the proteins of both subtypes**.

Format: XLSX, Size: 75Kb

**Additional file 2: This file contains the rule-based models for all the proteins of both subtypes**.

Format: XLSX, Size: 42Kb

**Additional file 3: This file contains list of names of the unseen viral sequences for both subtypes that were either miss-classified or could not be classified by the rule-based models**.

Format: XLSX, Size: 11Kb

**Additional file 4: This file contains singular and combinatorial signatures from the rules for both subtypes**.

Format: XLSX, Size: 75Kb

**Additional file 5: This file contains all the phylogeny trees marked with top 5 rules**. Each sequences is represented by its GeneBank accession. The nodes with a LightBlue background are the new, unseen sequences. The unmarked nodes do not support the top 5 rules, and were either supporting rules other than the top 5 or were not classified by the models.

